# ARHGEF3 Regulates Hair Follicle Morphogenesis

**DOI:** 10.1101/2024.09.13.612256

**Authors:** Krithika Kalyanakrishnan, Amy Beaudin, Alexandra Jetté, Sarah Ghezelbash, Diana Ioana Hotea, Jie Chen, Philippe Lefrançois, Mélanie Laurin

**Affiliations:** Centre de recherche du CHU de Québec – Université Laval, axe Oncologie, Québec, Canada; Programme de biologie moléculaire et cellulaire, Université Laval; Département de biologie moléculaire, biochimie médicale et pathologie, Université Laval; Faculté de médecine, Université Laval; Centre de recherche sur le cancer de l’Université Laval (CRC); Centre de recherche en organogénèse expérimentale (LOEX); Cancer Axis, Lady Davis Institute for Medical Research, Montreal, QC, Canada; Division of Experimental Medicine, McGill University, Québec, Canada; Department of Cell & Developmental Biology, University of Illinois at Urbana-Champaign, Urbana, IL, USA; Department of Biomedical and Translational Sciences, Carle Illinois College of Medicine, Urbana, IL 61801; Division of Dermatology, Department of Medicine, McGill University, Montréal, QC, Canada

**Author notes:** **Corresponding author’s email address:**.

**Keywords:** morphogenesis, hair follicle, ARHGEF3, placode, RhoGEF, Rho GTPase, P-cadherin

## Abstract

During embryogenesis, cells arrange into precise patterns that enable tissues and organs to develop specialized functions. Despite its critical importance, the molecular choreography behind these collective cellular behaviors remains elusive, posing a major challenge in developmental biology and limiting advances in regenerative medicine. By using the mouse hair follicle as a mini-organ system to study the formation of bud-like structures during embryonic development, our work uncovers a crucial role for the Rho GTPase regulator ARHGEF3 in hair follicle morphogenesis. We demonstrate that *Arhgef3* expression is upregulated at the onset of hair follicle placode formation. In *Arhgef3* knockout animals, we observed defects in placode compaction, leading to impaired hair follicle downgrowth. Through cell culture models, we show that ARHGEF3 promotes F-actin accumulation at the cell cortex and P-cadherin enrichment at cell-cell junctions. Collectively, our study identifies ARHGEF3 as a new regulator of cell shape rearrangements during hair placode morphogenesis, warranting further exploration of its role in other epithelial appendages that arise from similar developmental processes.

## INTRODUCTION

The molecular mechanisms that coordinate collective cell behaviors during organogenesis remain poorly understood. Hair follicles in mouse skin serve as ideal mini-organs for studying these processes due to their abundance, spatial pattern, and global alignment within the epidermal plane (1). During embryonic development, hair follicle progenitors are specified in the epidermis through paracrine and reciprocal signaling between the epidermal and underlying dermal compartments (2). Extensive loss- and gain-of-function experiments have elucidated the sequence of actions of key developmental pathways including BMP, WNT, SHH, and FGF that regulate hair follicle cell specification, downgrowth, and differentiation (3). More recent studies have also highlighted how mechanical forces influence and shape the architecture of the skin (4–9). Despite these significant advances, only a few downstream molecular effectors have been identified as crucial in mediating the effects of upstream signaling and mechanical input during hair follicle morphogenesis (10,11).

The development of the skin epidermis begins when cells from the surface ectoderm commit to an epidermal fate (3). By balancing proliferation and differentiation, these progenitors generate a stratified squamous epithelium, comprising an innermost proliferative basal cell layer and three differentiated keratinocyte layers, which are critical for the skin’s barrier function (12). Additionally, some of these progenitors give rise to skin appendages, such as the hair follicles (13). The first distinct morphological sign of hair follicle development is the formation of an epithelial thickening known as the hair placode. Directional cell migration and cell compaction have been shown to promote placode formation (14). Following the elongation of placode cells, external contractile forces in both the epidermal and dermal compartments cause these cells to expand their basal surface while the apical surface remains unchanged, which promotes their invagination (7). As remodeling of the extracellular matrix occurs around the placode, the pressure on placode cells is reduced, facilitating their reentry into mitosis (7). Further downgrowth of the bud is achieved through oriented cell divisions, leading to the formation of hair germs and hair pegs (7,15). Genetic manipulations, such as the loss of Myosin IIa (*Myh9* knockout mice) or treatment of mouse embryos with inhibitors of actomyosin remodeling, have highlighted the crucial role of cytoskeletal components in these processes (7,10,11,14,16,17).

In mouse skin, the alignment of hair follicles along the anterior-posterior axis of the embryo and the polarization of their downgrowth are governed by planar cell polarity (PCP), which refers to the coordinated polarization of a field of cells within a tissue plane (18,19). As the epidermis develops, significant changes occur in the shape and orientation of basal cells (10,20). These changes coincide with and are essential for the partitioning of PCP proteins, such as CELSR1, along the anterior-posterior axis of the basal cells. Disruption of epidermal contractility perturbs the establishment of PCP cues in the epidermis and leads to the misorientation of hair follicles (10,20). Importantly, mutations in conserved PCP components, such as *Frizzled-6*, *Celsr1* and *Vangl2,* result in misalignment of developing hair follicles (19,21–24). Still, the molecular effectors that act downstream of these core PCP components to coordinate cell rearrangements in the hair follicle remain to be identified. Understanding how PCP is established in skin is vital for comprehending how polarity is coordinated among neighboring cells and how it is manifested at tissue level.

In addition to the signals provided by epidermal cells, proper hair follicle morphogenesis requires extensive cellular rearrangements within the developing hair follicle. In the placode, counter-rotational cell movements play a crucial role in ensuring the polarization of the hair follicle and the asymmetric positioning of progenitor cells (16). These rearrangements are reminiscent of convergent extension movements, which promote directional elongation via cell intercalation and junctional shrinkage (16,25,26). While PCP components could help bias junctional contraction in one direction, another mechanism proposed for facilitating cell movement and neighbor exchange in the placode is the upregulation of the adherens junction component P-cadherin (also known as Cadherin-3) and the concurrent downregulation of E-cadherin in the central cells of the placode (27–29). Following rotational movements, P-cadherin-enriched cells become positioned in the anterior region of the polarized hair follicle, while E-Cadherin remains downregulated (16,19). This transition is crucial, as overexpression of E-cadherin can inhibit hair follicle formation by preventing invagination (27,30). Again, the molecular mechanisms downstream of these adherens junctions that contribute to hair follicle morphogenesis remain poorly understood.

Recently, we utilized our ability to transduce epidermal progenitors by injecting lentiviral particles into the amniotic cavities of mouse embryos using ultrasound guidance (31). This method allowed us to conduct an RNAi-mediated screen to identify new regulators of epidermal and hair follicle morphogenesis among components of the Rho GTPase networks, which are key cytoskeletal regulators (32–35). One of the candidates identified is ARHGEF3, also known as XPLN (36,37). Our findings revealed that cells transduced with shRNAs targeting ARHGEF3 failed to contribute to hair follicles, although their representation in the epidermis remained unchanged (32). This suggests that ARHGEF3 is a positive regulator of hair follicle development but is not essential for the formation of the epidermal barrier.

ARHGEF3 functions as a RhoGEF for RHOA and RHOB via its DH-PH GEF domain (36). In addition to its GEF activity, ARHGEF3 acts as a negative regulator of mTORC2, inhibiting signaling to AKT and restricting myoblast differentiation (38). In injured muscles, ARHGEF3 operates differently in a RHOA-dependent manner to restrict muscle regeneration. Remarkably, muscles in *Arhgef3^-/-^* knockout animals repair more effectively through the activation of autophagy (39). Another *Arhgef3*-null mouse model revealed that depletion of this RhoGEF leads to larger platelets without impairing their function (40). In a disease context, ARHGEF3 has been shown to promote the stability of ACLY, an ATP citrate lyase, by preventing its association with the E3 ligase NEDD4 in lung cancer cells (41). Despite these findings, the biological and molecular functions of ARHGEF3 remain largely unexplored. Here, we examine how ARHGEF3 regulates hair follicle morphogenesis by modulating cell compaction and P-cadherin-mediated cell-cell junctions.

## RESULTS

### ARHGEF3 is expressed in the developing skin

Of the 26 potential regulators of hair follicle morphogenesis identified in our screen, *Arhgef3* was the only one found to be differently expressed between the hair placodes and the interfollicular epidermis across multiple studies (29,32,42–45). Indeed, analysis of transcriptomic datasets from the *Hair-GEL* and *Sulic et al.* platforms revealed that *Arhgef3* mRNA levels were 2.57 times higher in the hair placodes compared to the epidermis (**Fig. 1A**) (44–46). Using these datasets, isoform switch analysis revealed that *Arhgef3* transcript variant 3, (*ENSMUST000000224981.2; NM_001289687.1*) was the most highly expressed isoform in the skin (**Fig. 1B**). Furthermore, the difference in gene expression between the placode and epidermis persisted when examining this specific transcript (**Fig. 1B**). To investigate whether this differential expression was observable directly in mouse skin tissue, we employed *in situ* hybridization on E18.5 mouse skin sections using a fluorescently labeled *Arhgef3* probe. Our results confirmed that *Arhgef3* mRNA levels were significantly higher in the hair placode compared to the epidermis, and this elevated expression of *Arhgef3* persisted in the developing hair follicle throughout its growth (**Fig. 1C**). This difference in expression between the two compartments was further appreciated when we compared it to the broad distribution of the ubiquitously expressed *Polr2a* mRNA in the same tissue (**Fig. 1C**). In summary, the upregulation of *Arhgef3* expression at the onset of hair follicle development suggests that it may play a crucial role during its morphogenesis.

**Figure 1.**
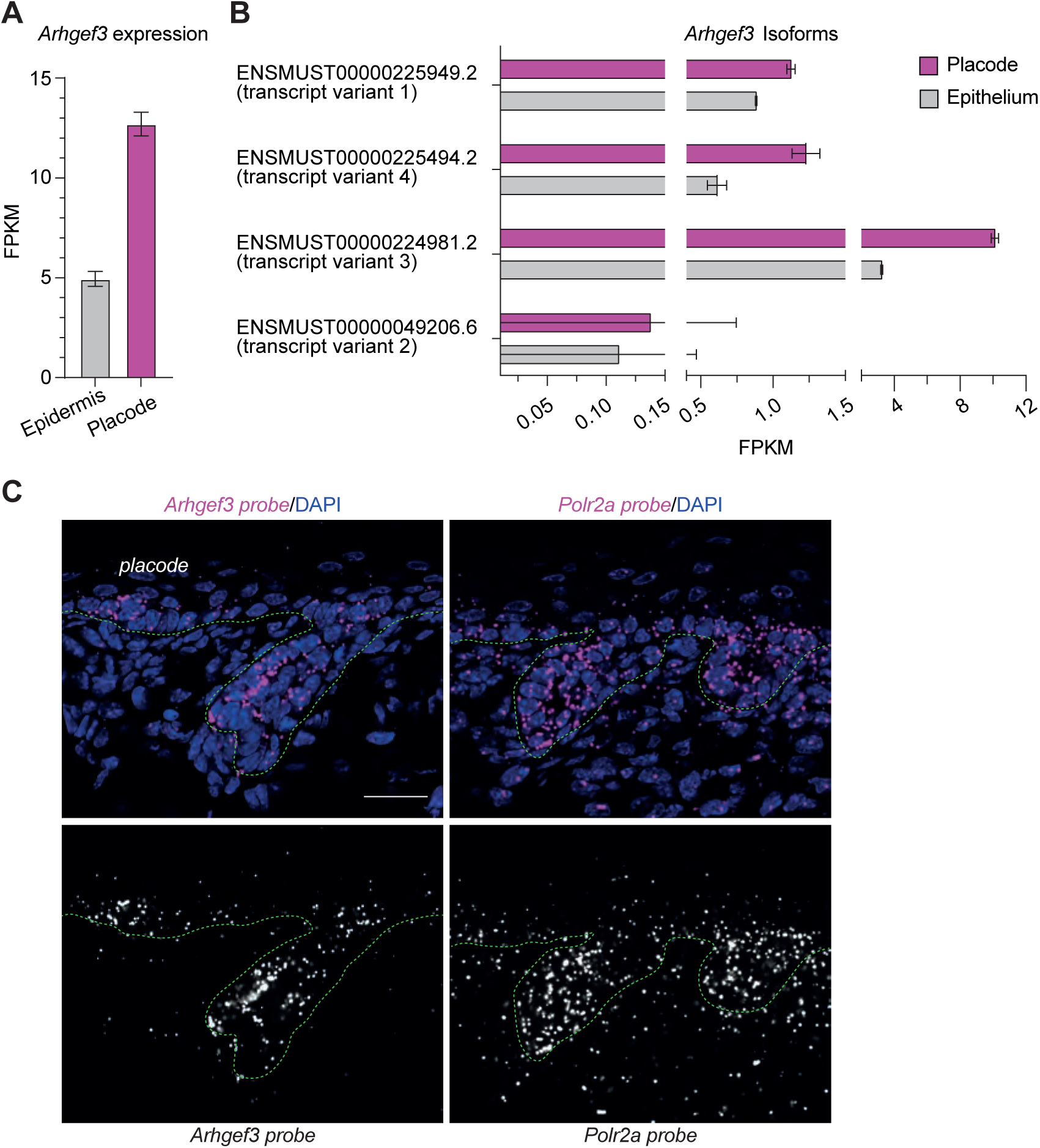
ARHGEF3 is expressed in the developing skin. (A) Gene-level RNA expression of *Arhgef3* in the hair placode and the epidermis using the *Hair-GEL* and *Sulic et al*. datasets, shown as mean fragments per kilobase per million (FPKMs) ± SEM (n=9). (B) Expression of *Arhgef3* transcript variants in the placode and the epidermis using the *Hair-GEL* and *Sulic et al.* datasets, shown as mean FPKMs ± SEM (n=9) (C) RNAscope® *in situ* hybridization (ISH) targeting *Arhgef3* mRNA (magenta) in the epidermis. Upregulation of transcript levels in the hair placode can be observed in the mouse skin tissue at E18.5 by the number of probe’s dots in this compartment. *Polr2a* mRNA is ubiquitously expressed in cells and used as a positive control. Scale bars: 25 µm. DAPI is used to label cell nuclei. Dotted green line is used to delimit the epidermis and hair follicles from the dermis.

### ARHGEF3 is not required for skin barrier formation

To examine the role of *Arhgef3* during skin development, we employed an *Arhgef3*-knockout mouse strain (*Arhgef3^-/-^*), generated by deleting a portion of exon 3, which is the first exon common to all four *Arhgef3* isoforms in mice (39). *Arhgef3-null* animals are viable, fertile, and exhibit enhanced muscle repair capabilities following injury, but their skin has not been thoroughly characterized (39). In our screen, the proportion of cells that expressed shRNAs targeting *Arhgef3* were not enriched or depleted in the epidermis at endpoint, which was similar to the behavior of cells expressing non-targeting shRNAs (32). This result suggests that ARHGEF3 is not crucial for epidermal development. However, because shRNA-mediated depletion can result in partial knockdown of their target and since ARHGEF3 has been shown to regulate the proliferation of other cell types (41,47), we investigated if the complete loss of ARHGEF3 in knockout embryos impairs epidermal proliferation. To explore this, pregnant females were pulsed with 5-ethynyl-2’-deoxyuridine (EdU) to label the cells in S-phase in the embryos. When quantifying the number of EdU^+^ basal cells, which are labelled by P-cadherin, we observed a slight decrease in their average number in *Arhgef3^-/-^* embryos compared to wild-type animals (**Fig. 2A, B**). Nevertheless, this difference was not statistically significant. Furthermore, measuring epidermal thickness using immunofluorescence for P-cadherin to identify the base of the epidermis revealed no significant differences between knockout and wild-type animals (**Fig. 2C, D**). Immunofluorescence analysis also revealed that embryos from both genotypes showed comparable levels of Keratin 10, a marker of differentiated suprabasal cells in the epidermis (**Fig. 2C**). Likewise, levels of LORICRIN and Filaggrin, predominantly expressed in the granular layer of the epidermis, remained unchanged in the absence of ARHGEF3 (**Fig. 2E, F**). Finally, we conducted a barrier assay from E16.5 to E18.5 in wild-type and *Arhgef3* knockout embryos and observed normal, development of the skin barrier in all animals (**Fig. 2G**). Collectively, these analyses indicate that ARHGEF3 is dispensable for epidermal development, which is consistent with the findings from our screen (32).

**Figure 2.**
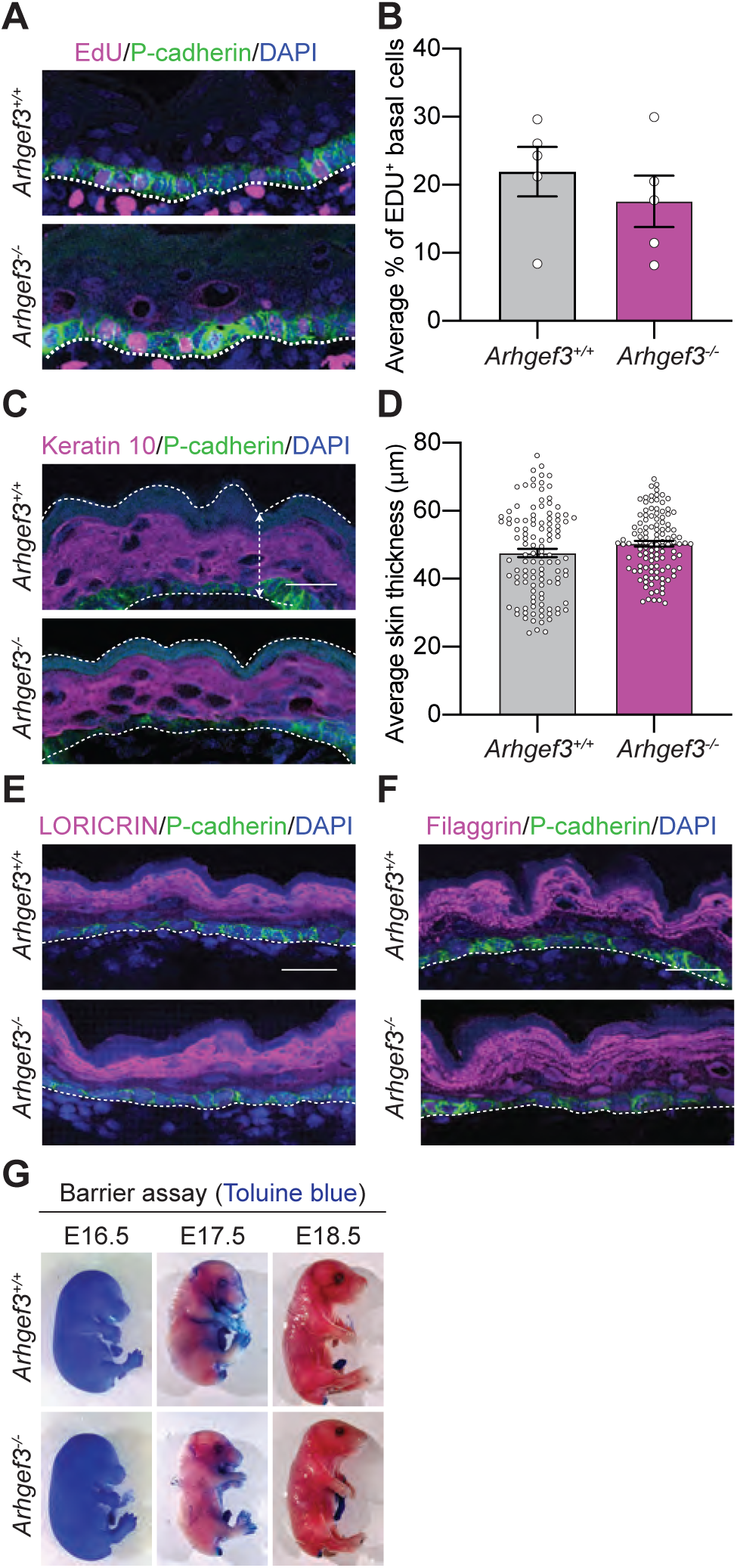
ARHGEF3 is not required for skin barrier formation. (A) EdU and P-cadherin immunofluorescence on E18.5 back skin sections. (B) Graph displays the quantification of the average percentage of basal (P-cadherin^+^) EdU^+^ cells ± SEM. Statistical analyses were performed using a two-tailed Mann-Whitney test, P value=0.4206, n=5 embryos per genotype from 5 litters and the experiment was independently performed 5 times. (C) P-cadherin and Keratin 10 immunofluorescence on E18.5 sagittal back skin sections. (D) Graph displays the average skin thickness ± SEM. Statistical analyses were performed using a two-tailed nested t-test, P value=0.5489, n=5 embryos (25, 30, 30, 15 and 15 measures) per genotype from 5 litters and the experiment was independently performed 5 times. (E) LORICRIN, (F) Filaggrin and P-cadherin immunofluorescence on E18.5 back skin sections. Representation of n=4 embryos. Proper expression of LORICRIN and Filaggrin is observed in both *Arhgef3^+/+^* and *Arhgef3^-/-^* animals. (G) Barrier assays on E16.5, E17.5 and E18.5 *Arhgef3^+/+^* and *Arhgef3^-/-^* mouse embryos. Representation of n=3. Scale bars: 25 µm. DAPI is used to label cell nuclei. Dotted white line is used to delimit the epidermis and hair follicle from the dermis.

### ARHGEF3 is required for hair follicle morphogenesis

With the confirmation that disrupting ARHGEF3 expression does not result in widespread epidermal defects, we proceeded to investigate whether this RhoGEF is essential for proper hair follicle development, as indicated by our screen (32). First, we assessed whether ARHGEF3 is required for the specification and initiation of hair follicle development. For this purpose, we used whole-mount immunofluorescence of P-cadherin on E16.5 back skin, which allowed us to visualize hair placodes, germs, and pegs from the staggered hair follicle waves (**Fig. 3A**). Quantitative analysis of these structures showed no significant difference in their average number between control and *Arhgef3*-null samples (**Fig. 3B**). To determine if the complete loss of ARHGEF3 affects cell proliferation in the hair follicle, we performed an EdU pulse experiment. Although there was a slight reduction in the number of EdU^+^ hair follicle cells in *Arhgef3^-/-^* embryos, this decrease was minimal and unlikely to have a significant impact on hair follicle morphogenesis. This was further supported by the quantification of hair peg length, which showed no significant difference between their average lengths in control and *Arhgef3^-/-^* embryos (**Fig. 3E**). Therefore, ARHGEF3 does not play a role in hair follicle specification and initiation.

**Figure 3.**
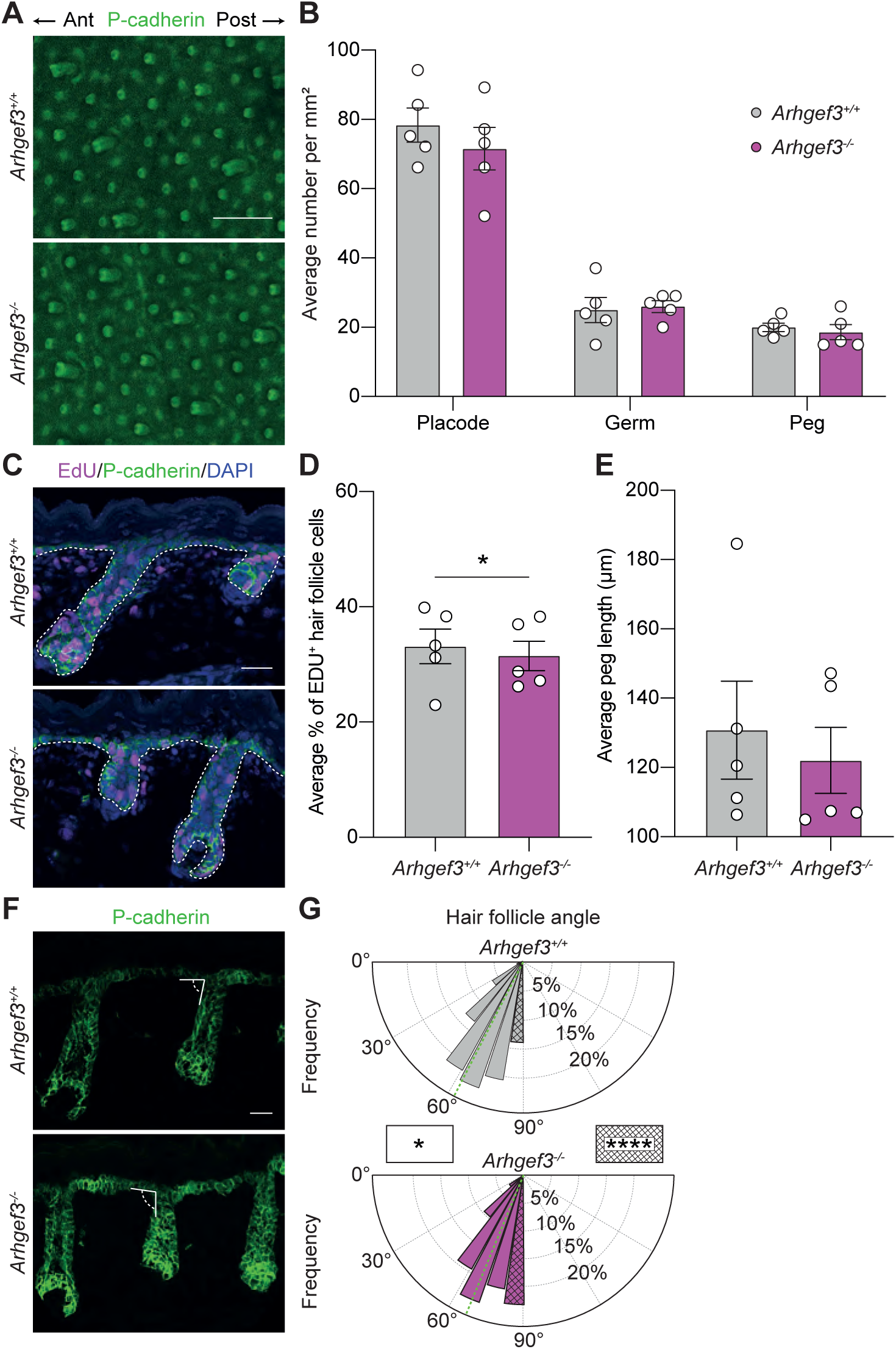
ARHGEF3 is required for proper hair follicle morphogenesis. (A) Maximum intensity Z-projections of P-cadherin whole-mount immunofluorescence of E16.5 back skin. Scale bars: 100 µm. The anterior – posterior axis of the embryo is indicated. (B) Graph displays the average number of hair placodes, germs, and pegs from the staggered hair follicle waves per 1 mm^2^ region of back skin ± SEM. Statistical analyses were performed using two-way ANOVA followed by multiple comparisons (with Sidak’s correction) between *Arhgef3^+/+^* and *Arhgef3^-/-^* conditions for each hair follicle stage. All comparisons were not significant (adjusted P values: placode = 0.5052; germ = 0.9967; peg = 0.9911), n=5 embryos per genotype from 5 litters and the experiment was independently performed 5 times. (C) EdU and P-cadherin immunofluorescence of E18.5 back skin sections. Scale bars: 25 µm. DAPI is used to label cell nuclei. Dotted white line is used to delimit the epidermis and hair follicle from the dermis. (D) Graph displays the average percentage of EdU^+^ hair follicle cells ± SEM. Statistical analyses were performed using a two-tailed Mann-Whitney test, P value=0.0312, n=5 embryos per genotype from 5 litters and the experiment was independently performed 5 times (hair follicles analyzed: *Arhgef3^+/+^*= 255; *Arhgef3^-/-^* = 205). (E) Graph displays the average length of hair follicle (pegs at E18.5) per embryo ± SEM. Statistical analyses were performed using a two-tailed nested t-test, P value=0.6359, n=5 embryos per genotype from 5 litters and the experiment was independently performed 5 times (peg analyzed : *Arhgef3^+/+^* = 23, 37, 19, 65, 71; *Arhgef3^-/-^* = 34, 39, 25, 52, 42). (F) P-cadherin immunofluorescence of E18.5 back skin. Highlighted is the strategy used to measure the angle between the basement membrane and developing hair follicle. Scale bars: 25 µm. (G) Rose plot displays the frequency of hair follicle angle (hair germ and peg) calculated in E with 10° bins. Dashed green line points out the circular mean (*Arhgef3^+/+^* = 62.77°; *Arhgef3^-/-^*= 67.40°). Statistical analyses to compare the distribution of angles were performed using Watson-Wheeler test for homogeneity of angles which, with many measurements, is approximately a Chi-square test with 2 degrees of freedom. This test reported 618 ties that were broken apart randomly. Hence, we proceeded to 20 iterations of the test that gave closely related P values: mean P value of iterations = 0.0208495±0.00023 (SD). To assess the difference in the frequency of straight hair follicles (striped bins,]80.0°-90.0°[) statistical analysis were performed using a two-sided Fisher’s exact test, P value<0.0001, n=5 embryos per genotype from 5 litters and the experiment was independently performed 5 times (hair follicles analyzed: *Arhgef3^+/+^*= 549; *Arhgef3^-/-^* = 545).

As hair follicles develop in mice, they typically align along the anterior-posterior axis of the embryo and penetrate the dermis at an angle relative to the basement membrane, rather than perpendicularly (19). In wild-type embryos, we measured that the hair follicles downgrowth within the dermis is evenly distributed around an average angle of 63 degrees relative to the basement membrane (**Fig. 3F, G** green line). Interestingly, *Arhgef3*-null hair follicles displayed a different distribution of angles around the increased average of 67 degrees (**Fig. 3G**; green line). In fact, we observed a significant increase in the percentage of straight-growing hair follicles in *Arhgef3^-/-^* embryos compared to the wild-type animals (**Fig. 3G**, dash bin in both genotypes), although their orientation along the anterior-posterior plane remained largely unchanged as observed in figure 3A. This resulted in a more entangled hair coat in shaved adult *Arhgef3^-/-^*animals, as the hair that fell away remained in clumps. This contrasted with the wild-type coat, in which individual hairs were easily dispersed.

PCP, which refers to the polarization of cells within the plane of an epithelium, is essential to the asymmetric downgrowth and alignment of hair follicles along the anterior-posterior plane of mice. As the skin develops, the progressive partitioning of core PCP components such as CELSR1 in epidermal cells provides instructive cues for hair follicle polarization (19). Using whole-mount immunofluorescence of CELSR1 and E-cadherin on E15.5 tissue, we investigated whether PCP is established in the absence of ARHGEF3, which could explain the defect in hair follicle downgrowth (**Fig. 4A**). First, we quantified the percentage of planar polarized cells, defined as cells with two opposing domains of CELSR1 at their cell surface membrane (10). On average, the percentage of these cells was similar in both *Arhgef3* knockout and wild type embryos (**Fig. 4A, B**). Quantification of the angle of polarization also showed that it was the same along the anterior-posterior axis of the embryos (**Fig. 4A, C**). These results suggest that CELSR1 domains are established in the epidermis of *Arhgef3* knockout animals and that the hair follicle angling defects observed in *Arhgef3*-null animals are likely uncoupled from the establishment of PCP in the epidermis.

**Figure 4.**
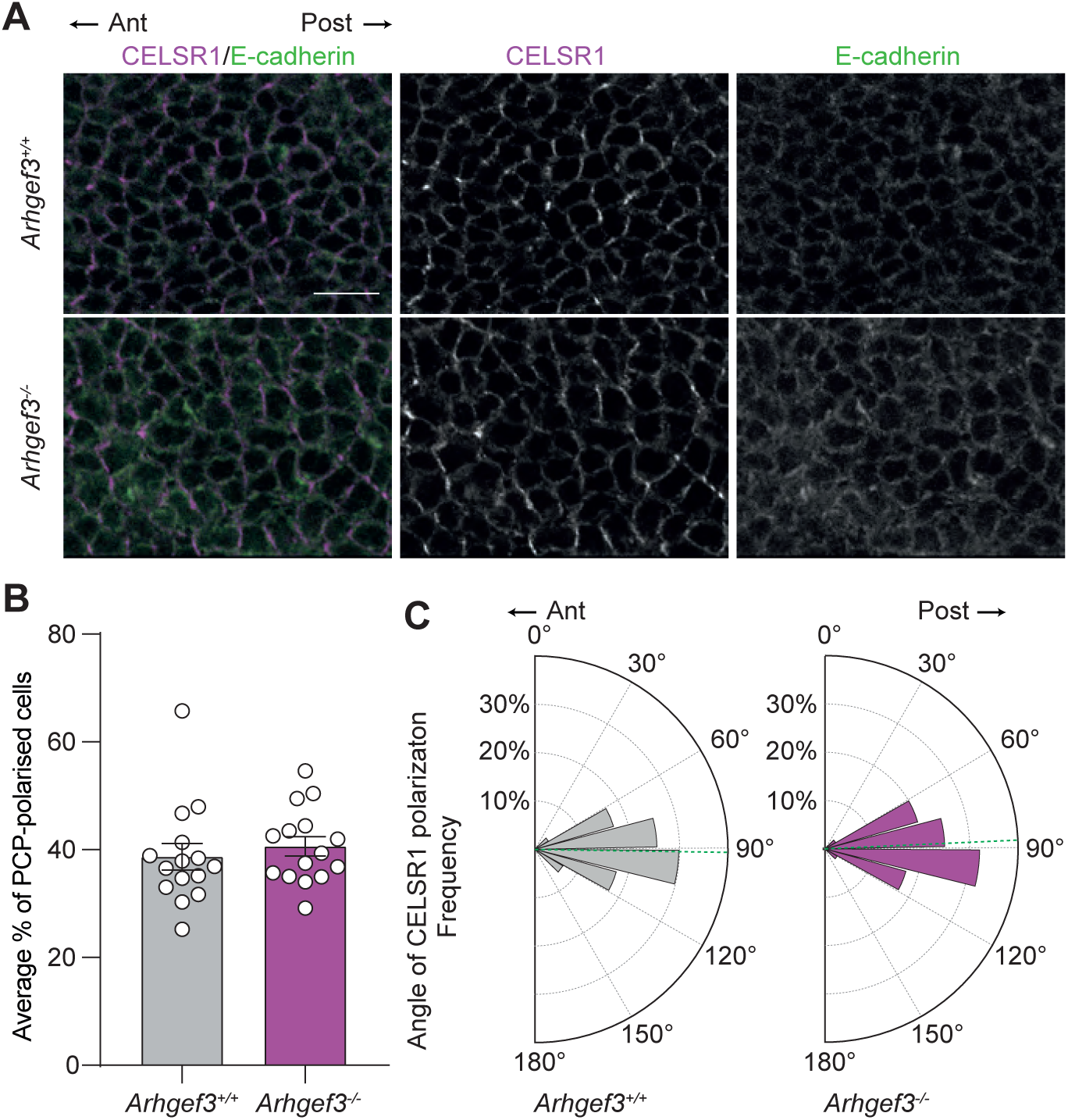
ARHGEF3 is not required to establish CELSR1 planar polarized domain in the epidermis. (A) CELSR1 and E-cadherin whole-mount immunofluorescence of E15.5 back skin. Scale bars: 25 µm. (B) Graph displays the average percentage of planar polarized cells ± SEM. Statistical analyses were performed using a two-sided Fisher’s exact test, P value=0.1403, n=3 embryos per genotype from 3 litters and the experiments were independently performed (cells analyzed: *Arhgef3^+/+^* =1473; *Arhgef3^-/-^* = 1450). (C) Rose plots display the orientation of the opposing CELSR1 domains in the epidermal cells, 90° being perfectly aligned with the anterior-posterior axis of the tissue (bins: 15°). Dashed green lines point out the circular mean (*Arhgef3^+/+^* = 90.91°; *Arhgef3^-/-^* = 88.97°). Statistical analyses to compare the distribution of angles were performed using Watson-Wheeler test for homogeneity of angles which, with a large number of measurements, is approximately a Chi-square test with 2 degrees of freedom. This test reported 646 ties that were broken apart randomly. Hence, we proceeded to 20 iterations of the test that gave closely related P values: mean P value of iterations = 0.7387±0.01286 (SD), n=3 embryos per genotype from 3 litters and the experiment was independently performed 3 times (cells analyzed: *Arhgef3^+/+^*=577; *Arhgef3^-/-^* = 540).

### ARHGEF3 regulates P-cadherin mediated junctions

Having established that CELSR1 domains are present in the epidermis at the onset of hair follicle development, we turned to a cell culture model of human keratinocytes (Ker-CT) to investigate the consequence of increasing the level of *Arhgef3* expression on cellular architecture and gain insights into the roots of the hair follicle defects. RT-qPCR analysis revealed that *ARHGEF3* mRNA is expressed in proliferating keratinocytes (Day 0) and remains consistently expressed throughout calcium-induced differentiation (Days 1 to 7) in culture (**Fig. 5A**). The differentiation of keratinocytes was confirmed by measuring the expression of *Keratin 10 (KRT10)* and *LORICRIN* mRNA, as both epidermis markers increase during differentiation (**Fig. 5A**). To mimic the increase in *Arhgef3* expression observed at the onset of placode formation, we generated a 3xFLAG-ARHGEF3 (3xF-ARHGEF3) doxycycline-inducible cell line by transducing a population of keratinocytes. Western blot analysis confirmed that the fusion protein was only expressed upon doxycycline exposure (**Fig. 5B**). Immunofluorescence showed that 3xF-ARHGEF3 localized to both the cytoplasm and nucleus in proliferating and differentiating keratinocytes (**Fig. 5C**). We then investigated the impact of increased ARHGEF3 levels on keratinocytes cell-cell junctions. In the presence of calcium, keratinocytes typically form E-cadherin-mediated cell-cell junctions that display a distinct honeycomb pattern. This pattern was observed in wild-type keratinocytes treated with doxycycline as well as in control 3xF-ARHGEF3 keratinocytes without doxycycline (**Fig. 5D**). When 3xF-ARHGEF3 was overexpressed, the effect on E-cadherin at the junction was modest. Although there was no significant enrichment or depletion of E-cadherin at the junction, ARHGEF3-overexpressing cells exhibited more continuous and less tortuous (*zipper-like*) E-cadherin staining at the cell-cell junction. Given that placode cells depend heavily on P-cadherin-mediated adherens junctions, we investigated its localization in our keratinocyte populations. Under control conditions, P-cadherin was evenly distributed at the cell-cell junctions between keratinocytes. Remarkably, we observed that an increase in ARHGEF3 expression was associated with a significant rise in P-cadherin levels at these junctions (**Fig. 5E**). This was particularly striking when looking at orthogonal views, which also revealed an increase in the cell height of ARHGEF3-overexpressing keratinocytes. To assess whether the observed rise in P-cadherin at the cell junctions was due to higher protein expression levels, we conducted Western blot analysis. Although both E-cadherin and P-cadherin levels increased during differentiation, no significant difference was detected between control and A ARHGEF3-overexpressing cells. This suggests that the strong recruitment of P-cadherin to cell junctions is not linked to an increased in its total protein levels (**Fig. 5F**). Therefore, ARHGEF3 appears to facilitate the relocalization of P-cadherin to cell-cell junctions in keratinocytes without altering its overall expression.

**Figure 5.**
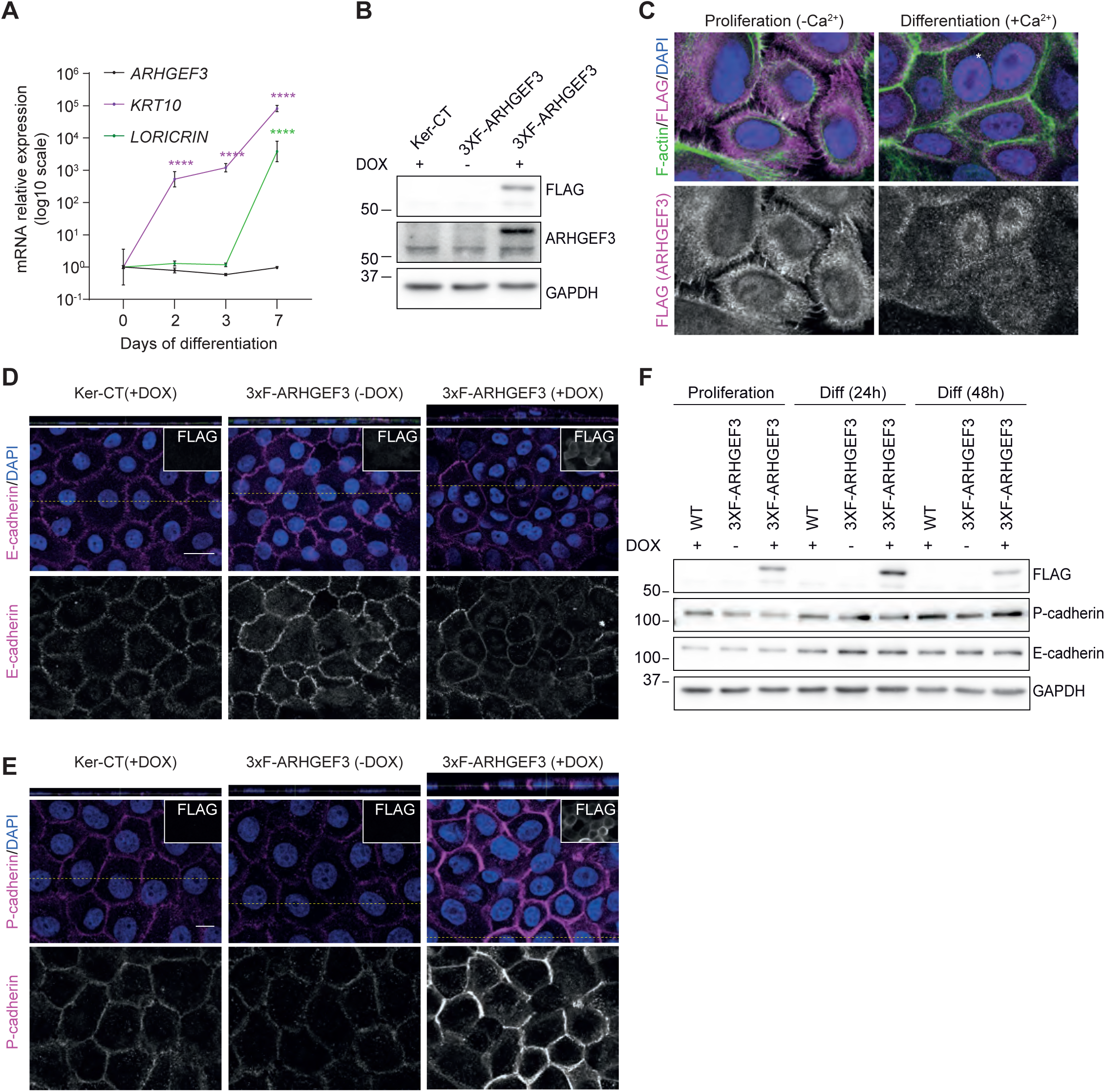
ARHGEF3 overexpression promotes the accumulation of P-cadherin at cell-cell junctions. (A) RT-qPCR analysis in proliferating (0) and differentiating Ker-CT (2, 3 and 7 days) showing the normalized expression of *ARHGEF3* mRNA on a log10 scale ± SEM. Proper differentiation of the keratinocytes was assessed using *KRT10* (Keratin 10) and *LORICRIN* mRNA expression (± SEM). Statistical analyses were performed using a two-way ANOVA followed by multiple comparisons with Dunnett’s correction to assess the difference between each differentiation time compared to proliferation state (D0), adjusted P value <0.0001, n=3 independent experiments, each with 3 technical replicates. (B) Western blot analysis for ARHGEF3, FLAG and GAPDH (loading control) showing the overexpression of 3xFLAG-ARHGEF3 in the transduced, doxycycline-treated cells. (C) FLAG and F-actin (Phalloidin) immunofluorescence on proliferating (without Ca^2+^; no cell-cell junctions) and differentiating (with Ca^2+^; with cell-cell junctions) keratinocytes. Scale bars: 10 µm. DAPI is used to label cell nuclei. (D) E-cadherin and FLAG immunofluorescence of keratinocytes grown for 48 hours in the presence of Ca^2+^. Scale bars: 10 µm. DAPI is used to label cell nuclei. Dotted yellow lines show the exact x slice where orthogonal views (y, z) were taken. (E) P-cadherin and FLAG immunofluorescence on keratinocytes grown for 24 hours in the presence of Ca^2+^. Scale bars: 10 µm. DAPI is used to label cell nuclei. (F) Western blot analysis for FLAG, E-cadherin, P-cadherin and GAPDH (loading control) in keratinocytes grown in proliferating and differentiating conditions with or without doxycycline as indicated.

### ARHGEF3 regulates placode compaction

The formation of cell-cell junction between keratinocytes is typically associated with the recruitment of F-actin at the junction and the formation of radial actin fibers, which could be observed in wild-type and control keratinocytes (**Fig. 6A**). However, ARHGEF3 overexpression caused a notable accumulation of F-actin at the cell cortex in keratinocytes, leading to cell compaction (**Fig. 6A**). This compaction was characterized by an increase in cell height, as seen in orthogonal views. Remarkably, in some clusters of ARHGEF3-overexpressing cells, keratinocytes appeared to stack on top of each other, a behavior not observed in control condition at this time point (**Fig. 6A**, right panel). Importantly, this multilayering was not due to premature differentiation, as these cells did not express the differentiation markers Keratin 10, 24 and 48 hours after the calcium switch (data not shown). This phenotype is strikingly similar to the cell compaction and cell elongation observed in hair placodes, which is driven by centripetal migration. To investigate whether defects in placode morphogenesis might explain the hair follicle phenotype observed in *Arhgef3^-/-^* animals, we analyzed placode formation using whole-mount immunofluorescence for P-cadherin on E16.5 back skin tissue. Consistent with previous studies, placode formation was associated with an increase in P-cadherin, in both wild-type and *Arhgef3-*null embryos (**Fig. 6B**). Since placode development is characterized by cell compaction, we measured the surface area of developing placodes using P-cadherin staining. Our measurements revealed that the average surface area of placodes was larger in *Arhgef3*-null animals (**Fig. 6C**). This suggests that increased *Arhgef3* expression at the onset of placode formation aids in the morphogenesis of placodes. Conversely, insufficient *Arhgef3* expression impairs hair follicle polarization, leading to a higher percentage of hair follicles growing straight.

**Figure 6.**
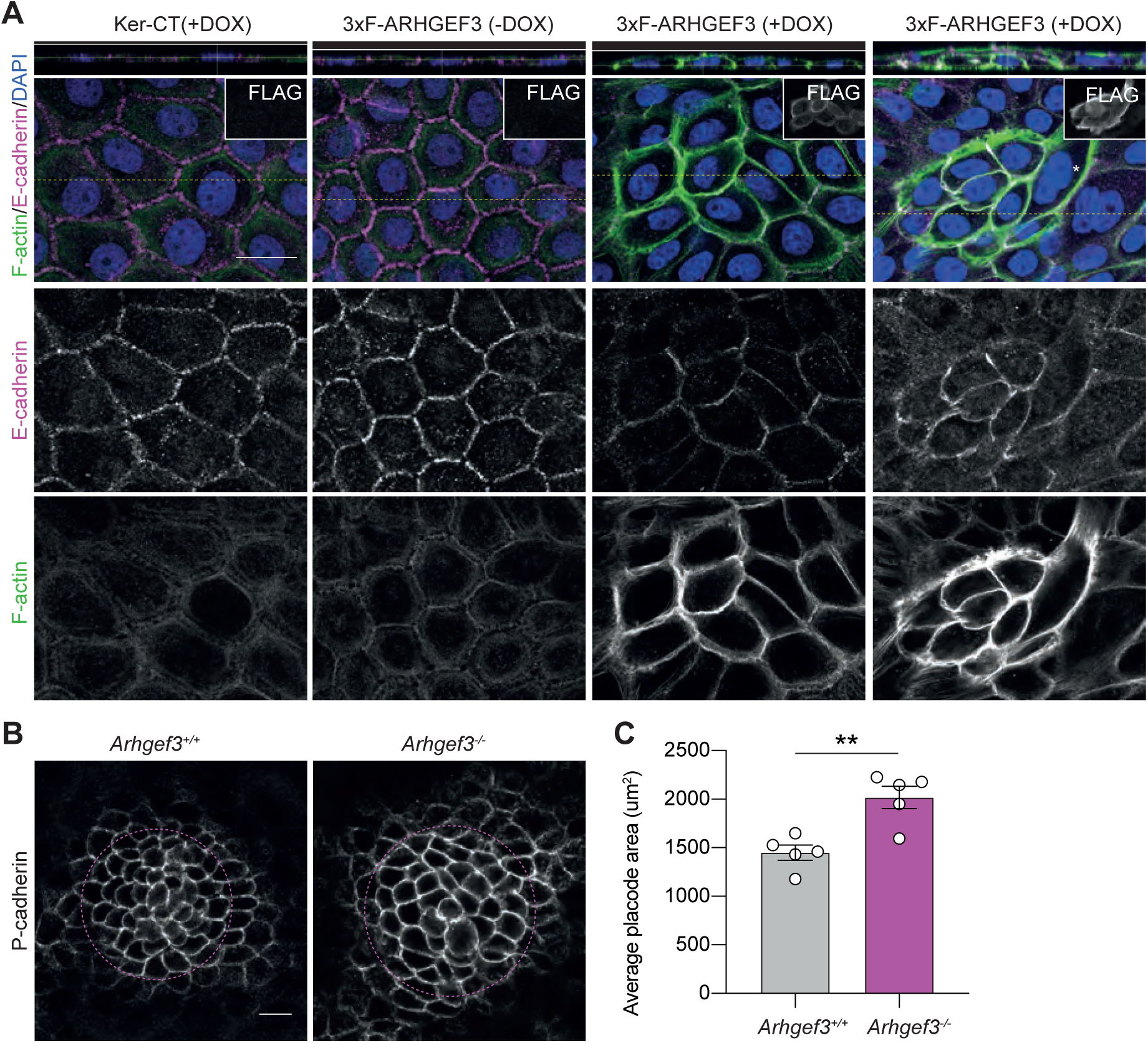
ARHGEF3 promotes placode compaction. (A) E-cadherin, FLAG, and F-actin (Phalloidin) immunofluorescence on keratinocytes grown for 24 hours in the presence of Ca^2+^ to allow cell-cell junctions to form. Scale bars: 10 µm. DAPI is used to label cell nuclei. Dotted yellow lines show the exact x slice where orthogonal views (y, z) were taken. (B) P-cadherin whole-mount immunofluorescence of E16.5 back skin. Scale bars: 10 µm. (C) Graph displays the average placode cell area ± SEM. Statistical analyses were performed using a two-tailed nested t-test, P value=0.0036, n=5 embryos per genotype from 5 litters and the experiment was independently performed 5 times (number of placodes analyzed: *Arhgef3^+/+^* = 43, 29, 25, 45, 30; *Arhgef3^-/-^* = 25, 48, 36, 44, 17).

## DISCUSSION

Building on our morphogenesis screen and transcriptomic data showing elevated ARHGEF3 expression in the placode relative to the epidermis, our study identified ARHGEF3 as a novel regulator of hair follicle morphogenesis (32,44,45). Although few studies have investigated ARHGEF3 regulation in mammalian cells, existing research shows that its expression is induced in muscles following their injury or in microglia after LPS treatment to mimic inflammation (39,48). Additionally, TGFß1 downregulates ARHGEF3, while HDAC inhibitors upregulate its expression in lung fibroblasts (41,49). Still, these findings provide limited clues into how ARHGEF3 is regulated during skin development.

Interestingly, studies on the Xenopus orthologue, *Arhgef3.2*, offer valuable insights. Arhgef3.2 plays a critical role in gastrulation, as demonstrated by loss-and gain-of-function experiments (50). In Xenopus, BMP4 restricts cell movement by inhibiting Arhgef3.2 transcription, while BMP inhibition promotes its expression (50,51). This is particularly intriguing given that BMP gradients in the epidermis also influence hair follicle formation—with high BMP levels inhibiting hair follicle development, and low BMP levels allowing placode initiation, which coincides with increased ARHGEF3 expression (2,3). This suggests a conserved regulatory mechanism across species, where an inverse relationship between BMP signaling and *Arhgef3* expression may be crucial for tissue morphogenesis.

The similarity between hair follicle morphogenesis and gastrulation is further underscored by the role of convergent extension cell movements in both processes. During Xenopus gastrulation, BMP inhibits convergent extension, while Arhgef3.2 regulates this process by interacting with Dsh2 via DAAM, two components of the non-canonical WNT/PCP signaling pathway, which is essential for convergent extension (50,52). Similarly, in hair follicle morphogenesis, PCP proteins are crucial not only for providing instructive cues from the epidermis but also for coordinating cellular movements within the developing hair follicle (16,19). These movements resemble convergent extension and are necessary for proper hair follicle polarization. While it remains unclear if ARHGEF3 functions downstream of Dsh2 in mammalian cells, our data suggest that ARHGEF3 acts either downstream or independently of core PCP cues in the skin, as these cues appear to be properly established in our system.

The collective behavior of cells depends heavily on the remodeling of cell-cell junctions. In hair follicles, this process is partly controlled by the upregulation of P-cadherin and the downregulation of E-cadherin (30,53). Our data suggest that while ARHGEF3 does not alter the overall levels of these adherens junction components in keratinocytes, increased ARHGEF3 levels significantly enhance P-cadherin accumulation at cell-cell junctions, with minimal impact on E-cadherin. Unraveling the molecular mechanisms that allow ARHGEF3 to selectively promote P-cadherin localization will be an exciting direction for future research. Currently, ARHGEF3’s known protein partners are limited, and expanding its interactome will likely deepen our understanding of its role in skin biology. Although ARHGEF3 exhibits both GEF-dependent and GEF-independent functions, its role in increasing F-actin accumulation at the cellular cortex of keratinocytes likely involves its RhoGEF activity. However, it remains unclear whether all ARHGEF3’s contributions to skin development depend solely on its ability to activate RHOA and RHOB.

While our research has primarily centered on ARHGEF3’s role in the epidermis and hair follicles, it is important to note that hair follicle polarization and progenitor cell asymmetry also rely on dermal contributions (7,16). Given that we employed a full knockout mouse model, the observed phenotypes could potentially be influenced by dermal signaling (39). However, no significant abnormalities were detected in the dermal cell population, and ARHGEF3 overexpression in keratinocytes resulted in cell-autonomous effects, such as F-actin accumulation at the cell cortex and P-cadherin relocalization at the membrane. To precisely address the role of dermal cells, developing an ARHGEF3 conditional mouse model would be invaluable and could help broaden our understanding of ARHGEF3 biological functions. Our findings highlight ARHGEF3 as a key regulator of hair follicle morphogenesis, linking it to conserved pathways governing placode formation and junctional remodeling via BMP gradients. Given the placode’s central role in various epithelial appendages, future studies will be essential to further explore ARHGEF3’s broader functions in these developmental processes.

## MATERIALS AND METHODS

### Animals models

*Arhgef3^-/-^* mice were previously generated and described (39). Animals were rederived upon their arrival at the CHU de Québec – Université Laval research center, where they are now maintained in a mouse-specific pathogen-free (SPF) facility. All mouse experiments were approved by Université Laval Animal Care Protection Committee, and they followed the Canadian Council of Animal Care Guidelines.

### RNA-Seq analysis

Whole-genome RNA-sequencing data of a total of 9 placode-enriched and 9 interfollicular epithelium (IFE) samples from E14.5 mice was collected from the *Hair-GEL* platform (2 placode-enriched and IFE sample pairs, Gene Expression Omnibus accession number GSE70288), and *Sulic et al*. (7 placode-enriched and IFE sample pairs, Gene Expression Omnibus accession number GSE212652). Raw fastq files were quality-checked with FastQC. Transcript-level quantification was obtained with Salmon (https://www.ncbi.nlm.nih.gov/pmc/articles/PMC5600148/) using selective alignment against the mm39 reference genome and transcriptome. Gene-and transcript-level fragments per kilobase per million (FPKMs) were extracted using the IsoformSwitchAnalyzeR R package (https://pubmed.ncbi.nlm.nih.gov/30989184/) and plotted using GraphPad Prism 10.

### RNAscope *in situ* hybridization

RNAscope *in situ* hybridization (ISH) was performed using the RNAscope Multiplex Fluorescent V2 Assay (Advanced Cell Diagnostics, 323270) according to the manufacturer’s protocol. Briefly, back skin from E18.5 embryo was dissected, fixed with 4% PFA for 1 hour at 4°C, washed several times with 1X PBS, and dehydrated in 20% sucrose overnight. The next day, the skin tissue was embedded and frozen in Tissue Plus O.C.T. Compound Clear (Fisher Scientific, 4585). Skin sections of 14 µm were generated, baked 30 minutes at 60°C, and fixed for 15 min at 4°C with 4% PFA. The sections were then dehydrated with serial incubations in increasing concentrations of ethanol (50%, 70%, 100% twice), treated with H_2_O_2_ for 10 minutes at room temperature, and with Protease IV (Advanced Cell Diagnostics, 322336) for 30 min at room temperature. Subsequent hybridizations (*Arhgef3* or *Polr2a* probes, 2 hours at 40°C) and amplifications (Amp1 (30 min, 40°C), Amp2 (30 min, 40°C) and Amp3 (15 min, 40°C)) were alternated with washes (twice, 2 min at room temperature) with 1X washing buffer (Advanced Cell Diagnostics, 310091). Both probes were in the C1 channel and fluorescence was developed using the HRP-C1 reagent, followed by TSA Vivid Fluorophore 570 (Advanced Cell Diagnostics, 323272; 1:2,000) and HRP blocker. Sections were counterstained with DAPI (Advanced Cell Diagnostics, 320858), mounted using Invitrogen^™^ ProLong^™^ Diamond Antifade Mounting media (ThermoFisher, P36970), and captured using an LSM-900 confocal microscope (Zeiss) with a LD C-Apochromat 40x water immersion objective (NA: 1.1). Basic image adjustments were performed in Fiji (ImageJ).

### Immunofluorescence, microscopy, and image processing

For whole-mount immunofluorescence on back skin tissues, embryos were fixed for 1 hour using 4% PFA at room temperature. The embryos were washed several times in 1X PBS while gently shaking and left to wash overnight. The following day the skin was dissected and blocked in gelatin buffer (1X PBS supplemented with 2.5% normal donkey serum (Sigma, 566460), 1% BSA (Wisent, 800-095-EG), 2% gelatin from cold water fish skin (Sigma, G7765) and 0.3% Triton X-100 (BioShop, TRX506.100) for at least 2 hours with agitation at room temperature. For whole-mount CELSR1 staining, the gelatin blocking buffer contained 2.5% fish gelatin, 2.5% normal donkey serum, 2.5% normal goat serum (Sigma, NSO2L), 0.5% BSA and 0.1% Triton X-100 in 1X PBS and the washes were done using 0.1% Triton X-100 in 1X PBS. Primary antibodies (see below) were diluted in gelatin buffer and incubated with agitation overnight at 4⁰C. The next day, the skin was washed 5 times with 0.3% Triton X-100 in 1X PBS. Secondary antibodies were diluted in gelatin buffer and incubated overnight with agitation at 4⁰C. The following day, the back skin was washed with 0.3% Triton X-100 in 1X PBS for more than 3 hours changing the solution at least 3 times, and then incubated with DAPI (Sigma, D9542; 0.2µg/ml) for 20 minutes. The nuclear stain was washed with 1X PBS followed by dH_2_O after which tissues were mounted on slides using Invitrogen^TM^ ProLong^TM^ Diamond Antifade Mounting media (ThermoFisher, P36970). Primary antibodies were used as follows: P-cadherin (Cadherin-3, R&D, AF761; 1:400), E-cadherin (R&D, AF748; 1:300) and CELSR1 (Fuchs lab gift, rabbit; 1:300). Secondary antibodies were used as follows: donkey anti-goat IgG cross-adsorbed Alexa Fluor 488 (ThermoFisher, A11055; 1:200) and donkey anti-rabbit IgG cross-adsorbed Alexa Fluor 594 (ThermoFisher, A21207; 1:200).

For immunofluorescence on cryosections, the back skin of embryos was fixed with 4% PFA for 1 hour at 4⁰C, washed several times with PBS 1X, and dehydrated in 20% sucrose overnight. The next day, the skin tissues were embedded and frozen in Tissue Plus O.C.T. Compound Clear (Fisher Scientific, 4585). Sections were fixed for 10 minutes in 4% PFA at room temperature, washed several times with PBS 1X, and blocked using gelatin buffer for 1 hour. Sections were incubated with primary antibodies diluted in gelatin buffer overnight at 4⁰C. The next day, sections were washed with 0.3% Triton X-100 and incubated with a secondary antibody (1:500) diluted in gelatin buffer for 1 hour. Later, these sections were washed with 0.3% Triton X-100, incubated with DAPI for 10 minutes and washed with PBS 1X. Sections were mounted using Invitrogen^™^ ProLong^™^ Diamond Antifade Mounting media. Primary antibodies were used as follows: P-cadherin (Cadherin-3, R&D, AF761; 1:400), Keratin 10 (BioLegend, 905403; 1:1,000), LORICRIN (BioLegend, 905104; 1:500) and FILAGGRIN (BioLegend, 905804; 1:400).

For immunofluorescence on Ker-CT, 1x10^5^ cells were plated in 15mm CultureWell^TM^ (ThermoFisher, C24776). Cells were fixed for 15 minutes using 4% PFA at room temperature and washed several times in 1X PBS. Next, cells were blocked in blocking buffer (1X PBS supplemented with 2.5% normal donkey serum, 1% BSA and 0.1% Triton X-100) for 1 hour with agitation at room temperature. Primary antibodies (see below) were diluted in blocking buffer and incubated overnight at 4⁰C. The next day, cells were washed once with 0.3% Triton X-100 in 1X PBS and twice with 1X PBS. Secondary antibodies (see below) were diluted in blocking buffer and incubated for 1 hour with agitation at room temperature. Cells were washed 3 times with 1X PBS and incubated for 10 minutes in 1X PBS containing DAPI. Cells were mounted on slides using Invitrogen^TM^ ProLong^TM^ Diamond Antifade Mounting media. Primary antibodies were used as follows: P-cadherin (Cadherin-3, R&D AF761; 1:400), E-cadherin (R&D AF748; 1:300) and FLAG-M2 (Sigma, F1804; 1:1,000). Secondary antibodies were used as follows: donkey anti-goat IgG cross-adsorbed Alexa Fluor (ThermoFisher, A11058 (594), A21447 (647); 1:1,000) and donkey anti-mouse IgG cross-adsorbed Alexa Fluor 488 (ThermoFisher, A-21202; 1:1,000). Phalloidin-iFluor 647 (Abcam, ab176759; 1:1,000) was used to label F-actin.

Images of whole-mounts, cryosections, and cells were captured using an LSM-900 confocal microscope (Zeiss) with either a Plan-Apochromat 20x air objective (NA: 0.8) or a LD C-Apochromat 40x water immersion objective (NA: 1.1). Basic image adjustments were performed in Fiji (ImageJ).

### Quantification of cell proliferation and hair follicle length

For cell proliferation assay, pregnant female mice were injected intraperitoneally with 5-ethynyl-2’-deoxyuridine (EdU, Sigma, 900584), allowing E18.5 embryos to be pulsed for 30 minutes. Embryos were dissected and the back skin was embedded in OCT as described above. EdU detection on cryosections was done according to the manufacturer’s instructions (Click-iT EdU Alexa Fluor 647 Imaging kit, Life Technologies, C10340). The ratio of EdU^+^ cells to all cells (DAPI^+^) was calculated for basal (based on P-cadherin^+^) and hair follicle (based on morphology) cells. To assess hair follicle length, the same cryosections were analyzed by drawing a line (straight or segmented) from the bottom of the basal layer until the end of the hair follicle and measured using Fiji (ImageJ).

### Quantification of hair follicle orientation and planar polarized cells

To assess hair follicle orientation, cryosections of E18.5 back skin were stained with P-cadherin to highlight the hair follicles and the basal layer in contact with the basement membrane. The angle between the basal layer and the hair follicle was drawn and calculated using Fiji angle tool as depicted in Figure 3. An angularity of more than 80° was classified as a straight hair follicle. To evaluate if PCP is established in the epidermis, CELSR1 and E-cadherin whole-mount immunofluorescence on E15.5 back skin was performed and 3 regions of 75 x 75 µm were analyzed per embryo. PCP-polarized epidermal cells were defined as cells in which opposing domains of CELSR1 were present. The frequency of the angle of these domains relative to the anterior-posterior axis of the embryo was determined using the Fiji straight-line tool.

### Barrier assay

Briefly, E16.5, E17.5 and E18.5 embryos were isolated from the pregnant mother. Euthanized embryos were immersed in ice-cold PBS 1X for 30 min. Embryos were immersed in a cold methanol gradient (1– 25%, 2–50%, 3–75%, 4–100% methanol) in water, and rehydrated in a methanol gradient in water (1–75%, 2–50%, 3–25%, 4–0% methanol), taking 2 minutes per step. Embryos were next immersed in 0.1% toluidine blue solution in water on ice for 2 minutes, with inversions. Embryos were destained in PBS 1X at least twice to reveal the dye pattern and barrier properties.

### Keratinocyte cell culture and cell differentiation assay

Ker-CT, an hTERT-immortalized keratinocyte cell line isolated from the foreskin of a male patient, was obtained from ATCC (CRL-4048). Cells were maintained as recommended in KGM^™^ Gold BulletKit™ media (Lonza, 192060) in a 37°C incubator in the presence of 5% CO_2_. For differentiation assay, 4x10^5^cells were plated in 9.6 cm^2^ wells. Differentiation was induced 24 hours following plating, when cells reached 80% confluency. For this, growth media was removed and replaced with KGM containing 1.5 mM of CaCl_2_ without growth factors. Total RNA was collected at 0-, 2-, 3- and 7-days following differentiation for further analyses.

### RNA isolation and RT-qPCR

RNA isolation was achieved using the Invitrogen™ PureLink™ RNA Mini kit (12-183-018A). Briefly, 1 µg of RNA was treated with DNase I (Thermo Scientific, EN0521) and retro-transcribed using the High-Capacity cDNA Reverse Transcription Kit (Applied Biosystems, 4368814). Semi-quantitative PCR was performed with the resulting cDNA using the LightCycler 480 SYBR Green I kit (Roche, 4707516001) and the primers listed below. The specificity and efficiency of primer pairs were defined prior to their usage.

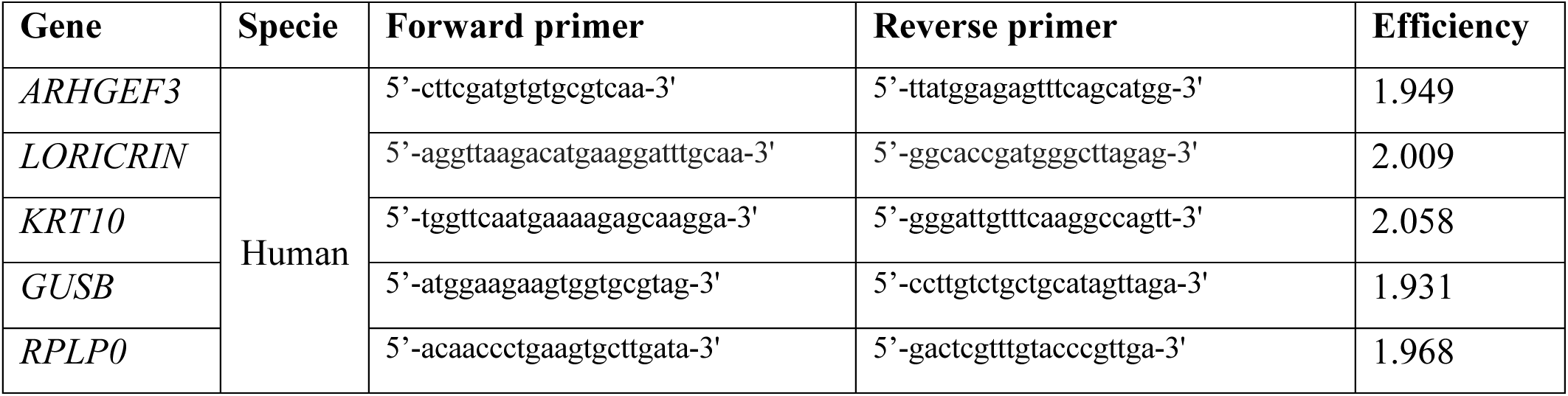

### Lentiviral production and infection

Envelope vector pPAX2 and packaging vector pVSV-G were kindly gifted by Amélie Fradet-Turcotte (Laval University, Quebec, Canada). The sequence of Arhgef3 isoform 3 with no ATG was cloned into a pCW57.1-3XFLAG pDEST vector. Lentiviral particles were produced by transfecting HEK293T cells with pPAX2, pVSV-G and 3XFLAG-ARHGEF3. Viral particles were collected after 48 hours of transfection and used to infect Ker-CT cells. Cells were selected with 1µg/ml puromycin and 3XFLAG-ARHGEF3 expression was induced using 50 ng/ml hygromycin B (BioBasic, BS725).

### Western blotting

Proteins were extracted from cells in RIPA lysis buffer containing 10 mM Tris-HCl pH 8.0, 1mM EDTA, 0.5mM EGTA, 1% Triton X-100, 0.1% sodium deoxycholate, 0.1% SDS, 140 mM NaCl, and supplemented with freshly added 1X proteases inhibitors (Roche, cOmplete EDTA-free Protease Inhibitor Cocktail, 11836170001). Lysates were centrifuged at 15,000 RPM for 15 minutes to remove debris. Samples were run on 8% polyacrylamide gel. Nitrocellulose membranes (Cytiva, 10600002) were incubated overnight with the following antibodies: FLAG (Sigma, F1804; 1:1,000), ARHGEF3 (ThermoFisher, PA5-30608; 1:5,000), Keratin 10 (BioLegend, 905403; 1:10,000), GAPDH (ThermoFisher, 39-8600; 1:10,000), E-cadherin (BD Biosciences, 610181; 1:500) and P-cadherin (R&D, AF761; 1:400). Secondary antibodies used were anti-mouse HRP (Millipore Sigma, A9044; 1:15,000), anti-rabbit HRP (Jackson Immunoresearch, 111-035-144; 1:15,000) and anti-goat HRP (Jackson Immunoresearch, 305-035-003; 1:15,000).

### Statistical Analysis

Statistical analyses were all performed with Prism 8 (GraphPad Software) unless stated otherwise. In all analysis, experiments were made independently for each pair of sibling embryos (*Arhgef3^+/+^* and *Arhgef3^-/-^*; sex as a biological variable was not considered given the embryonic nature of the analysis). For proliferation rate based on EdU^+^ cells for both basal layer and hair follicle analysis, the total number of EdU^+^ P-cadherin^+^ cells per embryo was normalized with the total number of P-cadherin^+^ cells (based on DAPI staining). The difference between proliferation rates was assessed with a two-tailed Mann-Whitney test with α=0.05 since there were only n=5 individuals per sample.

For both skin thickness, peg length and placode area, measurements were made multiple times on n=5 individuals per genotype and the difference between these two groups was assessed with a two-tailed nested t-test with α=0.05. Nested t-test permits to adjust the well-known t-test for the multiple measurements made in the same individual.

For the average number of hair placodes, germs, and pegs per 1 mm^2^ region of back skin, a two-way ANOVA was performed followed by multiple comparisons for each hair follicle stage with n=5 embryos per genotype. Sidak’s correction was applied to adjust for those multiple comparisons and adjusted P values are reported (with starting α=0.05).

For the relative expression of mRNA, a two-way ANOVA was performed on the transformed data of n=3 independent experiments (each with 3 technical replicates). Multiples comparisons were made with Dunnett’s correction to assess the difference between each differentiation time compared to proliferation state and adjusted P value are reported (with starting α=0.05).

For hair follicle angle as well as for CELSR1 angle, the circular mean was calculated. In brief, all individual angle measures (θ) were transformed to radians and then in a (x, y) format using this formula: (x, y) where x=sinθ and y=cosθ. The sum of those (x, y) coordinates was calculated and reverted to degrees to give the circular mean=deg(tan(Σ(x), Σ(y))). For both datasets, a Watson-Wheeler test was performed with R Studio (2024.04.2) to compare the distribution of the angles between *Arhgef3^+/+^*and *Arhgef3^-/-^* animals. This non-parametric test (for which the null hypothesis is that the two samples of angles come from the same population) was chosen to consider the fact that the angles measured could never be fully circular due to the nature of the measurement. When many measures are taken, the Watson-Wheeler test for homogeneity of angles is approximately a Chi-square test with 2 degrees of freedom. For hair follicle angle, a total of 10 embryos (5 per genotype) were analyzed from 5 independent experiments (hair follicles analyzed: *Arhgef3^+/+^* = 549; *Arhgef3^-/-^* = 545). For CELSR1 domains angle, a total of 6 embryos (3 per genotype) were analyzed from 3 independent experiments (cells analyzed: *Arhgef3^+/+^*=577; *Arhgef3^-/-^* = 540). All ties were randomly broken apart, giving a higher uncertainty on the P values obtained. To provide more information on that variability, we proceeded to 20 iterations of the test and reported the mean P value ± standard deviation (SD). We then performed a two-sided Fisher’s exact test on the same datasets to assess the difference in the number of straight hair follicle (]80.0°-90.0]) and the number of polarized cells (visually determined with opposing CELSR1 domains), respectively.

## ACKNOWLEDGEMENTS

We thank Steve Bilodeau for discussions; Patrick Laprise and Kathie Cockburn for the critical reading of the manuscript; Dr. Elaine Fuchs for the CELSR1 antibody; Carl St-Pierre from the “*Unité d’imagerie cellulaire, axe Oncologie – site de l’Hôtel-Dieu de Québec”* for assistance with microscopy; and the animal health technicians from the CHU de Québec – Université Laval-LOEX research center for mouse colony maintenance and animal care.

